# Multi-omics uncovers nutrient stress-driven interactions in the *Prymnesium parvum* holobiont, with vitamin B12 limitation highlighting mutualism

**DOI:** 10.64898/2026.01.22.701001

**Authors:** Lou Patron, Florian Petrilli, Anaelle Bout, Marinna Gaudin, Léna Gouhier, Clarisse Hubert, Damien Réveillon, Samuel Chaffron, Enora Briand, Matthieu Garnier

**Author notes:** Corresponding Authors Lou Patron, Enora Briand, Matthieu Garnier.

## Abstract

Microalgal-bacterial interactions are central to nutrient cycling and ecosystem functioning in marine habitats, yet the mechanisms structuring these associations under defined nutrient constraints remain poorly resolved. Using a synthetic 15-member bacterial community (SynCom) and controlled nitrogen (N), phosphorus (P), and vitamin B12 limitation, we investigated how nutrient scarcity shapes the physiology, metabolism, and transcriptional activity of the harmful alga *Prymnesium parvum* and its associated microbiota. Under N- and P-limitation, the SynCom had minimal impact on algal growth despite nutrient-dependent shifts in toxin production and metabolite profiles. In contrast, B12-limitation triggered a strong mutualistic interaction in which the SynCom enabled a three-fold increase in algal biomass and drove restructuring of intracellular and extracellular metabolomes, including the accumulation of ectoine and membrane-associated lipids and the depletion of thiamine-like and stress-associated metabolites. Metabarcoding revealed stable community composition but enrichment of B12-producing taxa under B12-limited conditions, while metatranscriptomics uncovered functional divergence among SynCom members. Bacteria segregated into specialized B12- or N-responsive strains and metabolically plastic generalists sustaining broad transcriptional activity across all nutrient regimes. B12 producers (*Marinovum algicola*, *Roseobacter* sp., *Halomonas* sp.) upregulated cobalamin biosynthesis exclusively under B12 limitation, whereas several dependent taxa induced B12 transport and B12-requiring enzymes, indicating active vitamin exchange within the holobiont. *P. parvum* displayed nutrient-specific transcriptional programs, with B12-limited co-cultures shifting toward growth-associated gene expression despite constitutive expression of the B12-dependent *metH* gene. These results demonstrate that vitamin auxotrophy acts as a key metabolic lever reorganizing holobiont function, driving reciprocal benefits and reprogramming algal-bacterial metabolism in our synthetic system.

## Introduction

Microalgae, as major constituents of phytoplankton, play an essential role in marine ecosystems by influencing biogeochemical processes at global scale. In natural environments, microalgae are associated with complex microbial consortia within the phycosphere, the microscale environment immediately surrounding algal cells [1] where symbiotic exchanges of metabolites can occur [2]. This web of interactions constitutes the algal holobiont, a functional ecological unit in which the algal host and its associated microorganisms interact metabolically and physiologically [3]. Reciprocal exchanges within the holobiont can range from mutualism to parasitism, collectively influencing the growth, nutrient acquisition, secondary metabolism, and community dynamics.

Nutrient limitations represent a major driver of microalgal physiology and holobiont dynamics. At global scale, nutrient availability, including macronutrients such as nitrogen and phosphorus, and micronutrients such as iron or vitamin B12 largely controls marine primary production [4]. At cellular level, nutrient limitation alters algal metabolism and growth. Nitrogen limitation redirects carbon fluxes toward storage compounds, notably lipids and carbohydrates [5–8] while phosphorus limitation impairs photosynthesis, triggers extensive lipid remodeling [9–12] and enhances alkaline phosphatases secretion [13]. Vitamin B12 limitation constrains growth in auxotrophic microalgae, primarily by blocking methionine metabolism [14]. Unlike macronutrient limitation, which triggers major metabolic reallocations, B12 deprivation mainly results in growth arrest, highlighting the reliance of many microalgae on bacterial cobalamin provision [15–18]. Yet, despite this well-established auxotrophy, the mechanisms governing how microalgae access, acquire, and compete for B12 with their associated microbial communities remain poorly understood [19].

Nutrient limitation also affects the bacterial partners within the phycosphere. Bacteria adjust their metabolism to scavenge limiting nitrogen and phosphorus, notably through high-affinity transporters and enzymes, like alkaline phosphatases [20, 21], and substitution of membrane phospholipids with non-phosphorus lipids [22, 23].

Indirectly, nutrient limitation alters algal physiology and exudation patterns, modifying the composition and lability of dissolved organic matter, thereby potentially selecting specific heterotrophic populations [24, 25]. Under B12 scarcity, algal growth is constrained, and bacteria capable of supplying the vitamin might gain a selective advantage, thus reshaping community composition [15, 26]. Overall, these interconnected responses illustrate that nutrient stress shapes both algal and bacterial physiology, making a holobiont-level approach essential to understand microalgal ecology from the micro-scale to ecosystem-scale.

Among the thousands of isolated microalgal species, 170 are known to produce toxins [27] and can form harmful algal blooms with major ecological and economic consequences [28]. In these species, nutrient availability influence not only growth and bloom dynamics but also toxin biosynthesis [29], making them highly informative models to connect environmental pressures, physiological and metabolic responses, and microbial interactions.

In this study, we use *Prymnesium parvum* as a model to explore how nutrient limitation shapes microalgal-bacterial interactions. *P. parvum* is a mixotrophic haptophyte with global distribution [30]. It is auxotrophic for vitamin B12 [31] and produces potent ichthyotoxins, the prymnesins, which have the potential to disrupt gill function in fish and have ecological and economic consequences worldwide [32–34]. Although prymnesin production increases under nitrogen and phosphorus stress [35], it remains unclear whether this response, whose underlying mechanisms remain to be elucidated, differs across nutrient limitations, and how bacterial communities influence growth and toxin production [36, 37].

Controlled holobiont models, including synthetic microbial communities (SynComs), offer a tractable alternative to study complex microbial interactions. SynCom-based studies have demonstrated that harmful algae can selectively enrich particular bacterial taxa [36], and that bacteria can protect algae from pathogens [38]. However, most SynCom studies focus on taxonomic or functional responses alone, while multi-omics approaches combining both algal and bacterial partners remain rare.

Here, we extend SynCom-based approaches to investigate *P. parvum*-bacteria interactions under nitrogen, phosphorus, and vitamin B12 limitation. By combining growth monitoring with multi-omics techniques including metabarcoding, metatranscriptomics, and metabolomics, we investigated both the taxonomic and functional responses of algal and bacterial partners. This holistic framework provides insights into how nutrient stresses shape *P. parvum* holobiont assemblage, and the extent to which bacterial communities modulate algal physiology, metabolism, and toxin content. Specifically, we test whether nutrient-specific bacterial assemblages drive distinct transcriptional and metabolic responses in *P. parvum*, with consequences for growth and toxin production. Through this approach, we aim to advance towards a mechanistic understanding of nutrient-dependent processes that underpin both algal ecology and biogeochemical functioning at larger scales.

## Material and methods

### Co-culture Experimental Design

Fifteen bacterial strains were selected for their ecological relevance and functional diversity, including vitamin B12 production and diazotrophy (Table 1), and assembled into the synthetic community (SynCom) as described previously [36]. *P. parvum* (CCAP 646/6) was acclimated for two weeks in L1 medium [39] under nitrogen (N-lim; N adjusted to 62 µM), phosphorus (P-lim; P adjusted to 2.2 µM), or vitamin B12-limited (B12-lim; B12 adjusted to 7.4 × 10⁻⁶ µM) conditions.

**Table 1.** Composition of the synthetic microbial community (SynCom) used in this study. The table lists the 15 bacterial strains, their taxonomic affiliation (phylum or class), and a functional trait of interest (vitamin B12 production or diazotrophy).

| Phylum/Class | Species | Functional traits |
| --- | --- | --- |
| **Bacteroidota** | *Cyclobacterium* sp. |  |
|  | *Maribacter* sp. |  |
|  | *Muricauda* sp. |  |
| **Alphaproteobacteria** | *Marinovum algicola* | B12 producer |
|  | *Marivita cryptomonadis* |  |
|  | *Roseobacter* sp. | B12 producer |
|  | *Thalassospira* sp. | B12 producer |
| **Gammaproteobacteria** | *Alteromonas* sp. 002 |  |
|  | *Alteromonas* sp. 005 |  |
|  | *Halomonas alkaliphila* | B12 producer |
|  | *Halomonas* sp. | B12 producer |
|  | *Marinobacter* sp. |  |
|  | *Vibrio aestuarianus* |  |
|  | *Vibrio diazotrophicus* | Diazotroph |
|  | *Pseudomonas stutzeri* | Diazotroph |

Cultures were inoculated at 1×10⁵ cells mL⁻¹ ± SynCom at the same concentration and grown in photobioreactors under continuous light (100 µmol photons m⁻² s⁻¹), at 20 °C, and pH 8.2, in a phenotyping bench as previously described [40]. Four biological replicates with SynCom and three without were monitored for 14 days and ecophysiological parameters were measured as described previously [36].

### Growth and nutrient monitoring

Algal and bacterial abundances were monitored daily respectively using a Multisizer 4 Coulter Counter (Beckman Coulter, Indiana, USA), and by flow cytometry (MACSQuant Analyzer 10, Miltenyi Biotec, Germany) after SYBR Green I staining (1%).

Nutrient concentrations (N, P, C) were measured on days 0, 3, 7, 9, and 14. For each photobioreactor, 15 mL of culture was filtered through pre-combusted GF/C filters, filtrates were used to quantify dissolved inorganic nitrogen and phosphate, and filters were kept for particulate organic carbon (POC) and nitrogen (PON) analyses. Dissolved N and P were measured on a Seal AutoAnalyzer (SEAL Analytical, Germany) using classical colorimetric methods [41] adapted for marine samples [42]. POC and PON were quantified by elemental analyzer (Flash 2000, Thermo Fisher Scientific, USA).

To confirm the limiting nutrient at the end of the experiment, cultures were individually supplemented with N, P, or vitamin B12 and growth recovery was monitored.

### 16S metabarcoding

16S-metabarcoding was performed as previously described [36] on the Syncom at day 0 and in the cultures at day 3, 7, 10 and 14 of the experiment. Briefly, samples were sequentially filtered through polycarbonate membranes of 3 µm and 0.2 µm pore size to separate *P. parvum*-associated bacteria (PA) from free-living bacteria (FL). DNA was extracted (NucleoSpin Plant II, Macherey-Nagel, Germany), amplified (16S, V3-V4 region, primers 301F (CCTAYGGGRBGCASCAG)/805R (GGACTACNNGGGTATCTAAT)) and sequenced on Illumina (ADNid, France). Sequences were processed using the SAMBA pipeline (https://gitlab.ifremer.fr/bioinfo/workflows/samba; v4.0.0), with taxonomic assignment of ASVs performed using SILVA (v138.1) as reference. Statistical analyses were conducted in R (v4.4.2) as described in section Statistical analyses. The sequencing dataset was deposited in the European Nucleotide Archive (ENA) under the project number PRJEB106332.

### Metabolomic and toxin analyses

Intracellular and extracellular prymnesins and metabolites were analyzed on day 8. Cells and supernatants were separated by centrifugation (50mL, 3,000 × g, 5 min at 4°C). Pellets were extracted twice with 1 mL of methanol (>99.9 % purity, CHROMASOLV™, Honeywell, USA), in an ultrasonic bath (15 min at 25 kHz). The two extracts were pooled, a subsample was ultrafiltered (0.2 µm, Nanosep MF, Pall, USA) and transferred into a glass vial while supernatants were processed by solid-phase extraction (SPE) using a Bond Elut C8 cartridge (500 mg, 10 mL, LRC, Agilent, USA), as previously described [43].

Data were acquired on a UHPLC system (1290 Infinity II, Agilent, USA) coupled to a high-resolution quadrupole time-of-flight mass spectrometer (Q-Tof 6550 iFunnel, Agilent, USA), and processed as described previously for prymnesins [43] and for metabolomics [36]. For the features significantly affected, MS/MS spectra were acquired by targeted MS/MS, at three collision energies (10, 30 and 50 eV). Spectra were processed using GNPS [44], SIRIUS v6.1.1 [45], and Flash entropy search [46] with the database retrieved as previously described [44]. Annotation levels ranged from 5 (exact mass) to 2 (probable structure by comparison with literature/library spectrum) [45]. All metabolomic data (full scan, MS/MS) are available at https://doi.org/10.12770/ as well as annotation details in Table S2.

### Transcriptomic Analysis

Metatranscriptomic analyses were performed on day 14. Cultures were centrifuged to obtain algal- and bacterial-enriched fractions (50 mL, 500 × g then 4,000 × g (10 min each)). RNAs from the bacterial enriched fractions were extracted (TRIzol-chloroform), precipitated (isopropanol), treated with DNAse I, quantified (Qubit fluorometer, Thermofisher Scientific, USA), rRNA-depleted (QIAseq FastSelect kit, Qiagen, Netherlands) and RNA integrity was assessed with a Bioanalyzer (Agilent, USA). Three samples per condition were sequenced (Illumina, Integragen, France), yielding 37-44 M paired-end reads per sample. Reads were trimmed (Cutadapt v4.1 [46]), residual rRNA removed (Bowtie2 v2.5.4 [47]) and mapped (HISAT2 v2.2.1 [48]) to the *P. parvum* genome (https://doi.org/10.12770/) and to the 15 bacterial genomes sequenced in this study (Table S1). Details about reads distribution as well as the taxonomic composition of SynCom mRNA reads are respectively available in Figure S1 and S2. Annotations of bacterial genomes were generated by merging the outputs of Bakta v1.11.4 [49], Prokka (v1.14.5) [50] and MaGe (v3.17.5) [51], all run with default parameters to reduce pipeline-specific biases. Count matrices for *P. parvum* and bacterial community genes were analyzed using DESeq2 (v1.46.0) [52] in R (https://www.r-project.org/) (v4.4.2). The sequencing dataset was deposited in the European Nucleotide Archive (ENA) under the project number PRJEB106332.

### Statistical analyses

Differences in prymnesin quotas, and cellular C/N ratios were assessed by a one-way ANOVA with Tukey’s HSD. Metabolomic data were log-transformed and Pareto-scaled prior statistical analyses using Metaboanalyst 6.0 (www.metaboanalyst.ca). For intracellular data, the matrices were additionally normalized by cell number before analysis. The fourth replicate B12lim with SynCom was identified as an outlier and excluded from all subsequent analyses. Principal component analysis (PCA), volcano plots (|log₂FC| > 2, padj < 0.05) and heatmaps were generated on the processed datasets.

Amplicon sequencing data were normalized using cumulative-sum scaling (CSS) in SAMBA. Community variation across nutrient conditions, fractions (PA vs FL) and time points was explored using PCA and Bray-Curtis dissimilarities. Differences in community composition were tested by PERMANOVA (adonis2, vegan).

For *P. parvum* transcriptomes, ribosomal protein genes were removed, raw counts normalized (median-of-ratios), and DEGs identified using Wald statistics with Benjamini-Hochberg correction (|log₂FC| > 3, padj < 0.05). Log₂-transformed counts were clustered using pheatmap.

For bacterial transcriptomes, high sparsity prevented direct DESeq2 application. A three-step gene filter was applied: detection in ≥2 replicates, mean expression >10 within a condition, and both criteria satisfied in ≥2 nutrient conditions; genes meeting the first two criteria in only one condition were labelled “ON” and defined as being exclusively expressed in that condition. Filtered genes were analyzed with DESeq2 using the same statistical thresholds as for *P. parvum*. Functional enrichment of bacterial DEGs was performed with clusterProfiler.

All statistics and figures (except for metabolomic PCA and metabarcoding barplots) are available on gitlab (https://gitlab.ifremer.fr/lp7b2b6/horus/)

## Results

### Growth monitoring

*P. parvum* was grown under N-, P-, and B12-lim (n=4 with SynCom; n=3 without SynCom). Cultures entered stationary phase at day 7 in all conditions. However, in B12-lim + SynCom, growth resumed on day 10, ultimately reaching a biomass 3.5 times higher at day 14. (Figure 1). In B12-Lim, algal cells sizes were significantly larger without SynCom (Figure S3). This SynCom-dependent recovery suggests with active alleviation of B12 limitation through microbial provisioning. Nutrient-spiking assays at day 15 confirmed successful establishment of the intended limitation (Figure S4-6). Bacterial dynamics were similar under N- and P-lim ± SynCom and B12-lim without SynCom. Concentrations were consistently higher with SynCom (1.6-fold in N-lim; 1.3-fold in P-lim). Bacterial growth of the SynCom was significantly enhanced in B12-lim, from the third day onwards ultimately reaching levels about 5-fold higher than under N-lim and 2-fold higher than under P-lim. These increases suggest reciprocal benefits in which algal exudates support bacterial proliferation under B12-lim.

**Figure 1.**
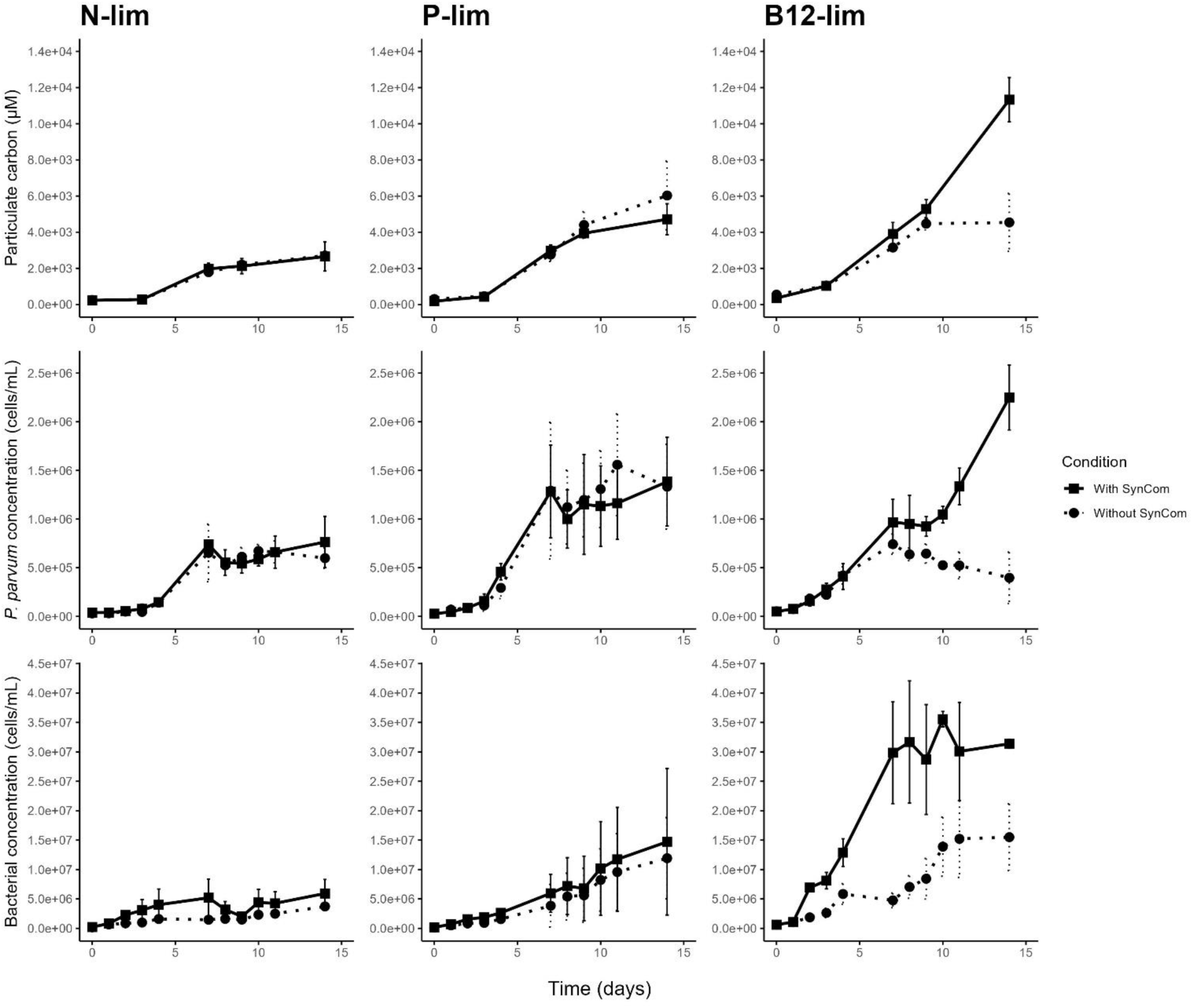
Growth dynamics of P. parvum and bacteria under nitrogen (N-lim), phosphorus (P-lim), and vitamin B12 (B12-lim) limitation, with and without a synthetic bacterial community (SynCom). Time-series of particulate organic carbon in µM (top row), algal cell concentration in cells/mL (middle row), and bacterial cell concentration in cells/mL (bottom row) in *P. parvum* cultures grown under nitrogen (left column), phosphorus (middle column), and vitamin B12 (right column) limitation. For each condition, cultures were supplemented (with SynCom, solid lines) or not (without SynCom, dashed lines) with a defined synthetic bacterial community at the beginning of the experiment.

### Toxin analysis

Intracellular and extracellular prymnesins (PRM) were quantified on day 8, when *P. parvum* was in a comparable physiological state across treatments. Intracellular PRM levels were highest under P-lim (1.5-fold higher than N-lim; 4-6-fold higher than B12-lim), while N-lim cultures showed 3-4-fold higher levels than B12-lim. The SynCom did not significantly affect PRM content in any nutrient condition (Figure 2).

**Figure 2.**
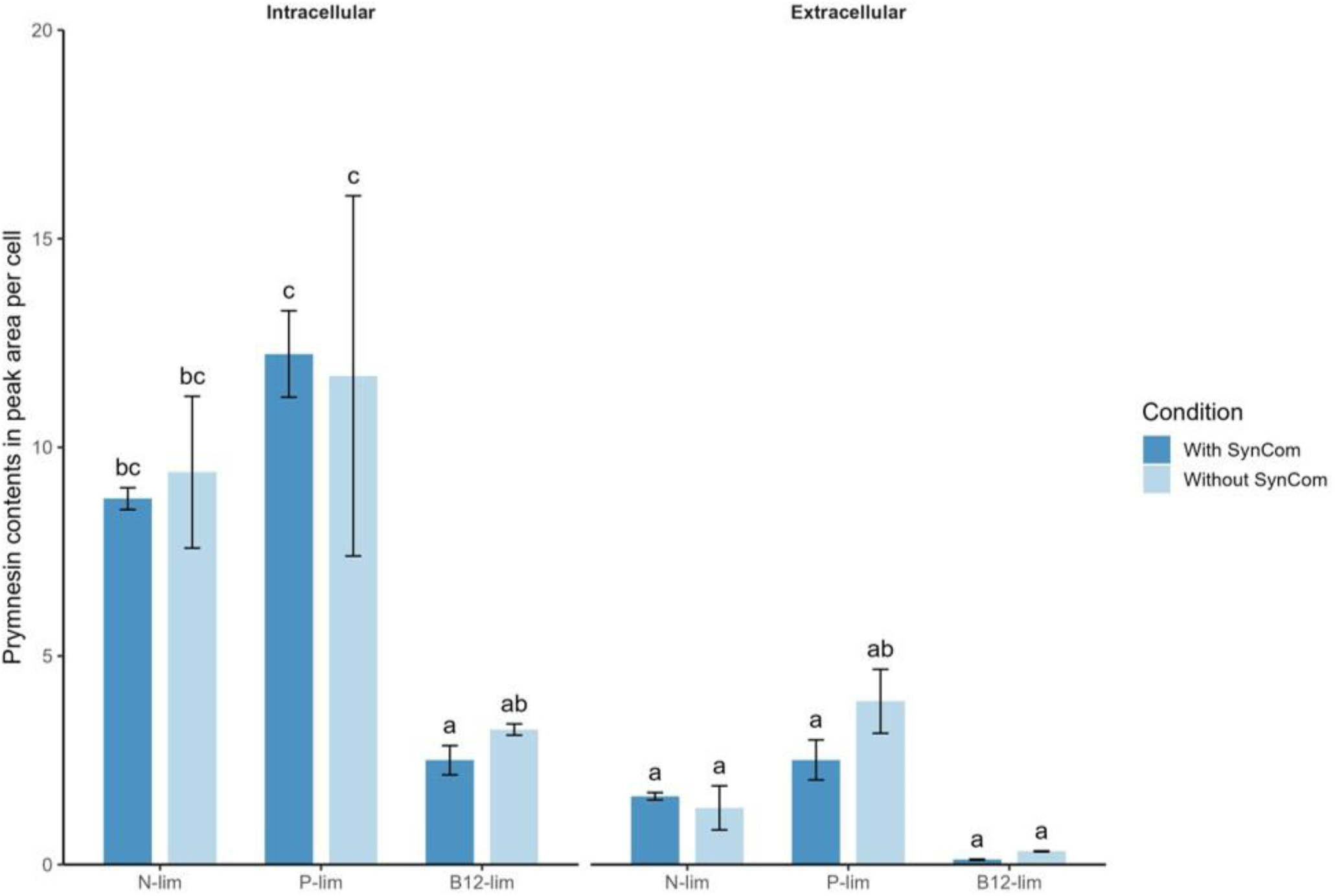
Intra- and extracellular prymnesin contents under different nutrient-limited conditions. Barplots show prymnesin levels (sum of PRM1 and 2, expressed as peak area per cell, mean ± SD) measured in N-lim, P-lim, and B12-lim *P. parvum* cultures, either with or without the presence of the SynCom. Bars are grouped by prymnesin fraction (intracellular vs extracellular) for each nutrient-limited condition. Measurements were performed at day 8 of the experiment.

### Metabolomic analysis

Metabolomic profiles revealed strong nutrient-dependent responses, for both intra- and extracellular fractions clearly clustered by nutrient conditions (Figure 3A and 3B), with a marked effect of the SynCom presence extracellularly under B12-lim (Figure 3B). This nutrient-driven structuring indicates distinct stress-specific metabolic responses, with B12-lim modulated by the SynCom. Comparisons of ±SynCom cultures highlighted 112 significantly modulated features including 72 intracellular and 40 extracellular in B12-lim, 6 extracellular in N-Lim and 7 extracellular in P-lim (Table S2). Among those systematically affected by the presence of the SynCom, we identified several extracellular and intracellular sulfobacins as well as intracellular α-amino-acid and benzene derivatives, hydroxysteroids and several unknown features. Among those affected by the presence of the SynCom in B12-lim only, with SynCom specific features, distinct intracellular and extracellular metabolic changes were observed. Intracellularly, DGCC-like lipids, PE-like lipids and one nitroaromatic compound were depleted, while ectoine and multiple PC- and PE-like lipids accumulated. Extracellularly, thiamine-like compounds, glycerolipids, steroids and glycosylated metabolites were reduced, whereas conjugated fatty acids, thiamine and the unknown feature M251T499 were strongly enriched (Figure 3C and 3D).

**Figure 3.**
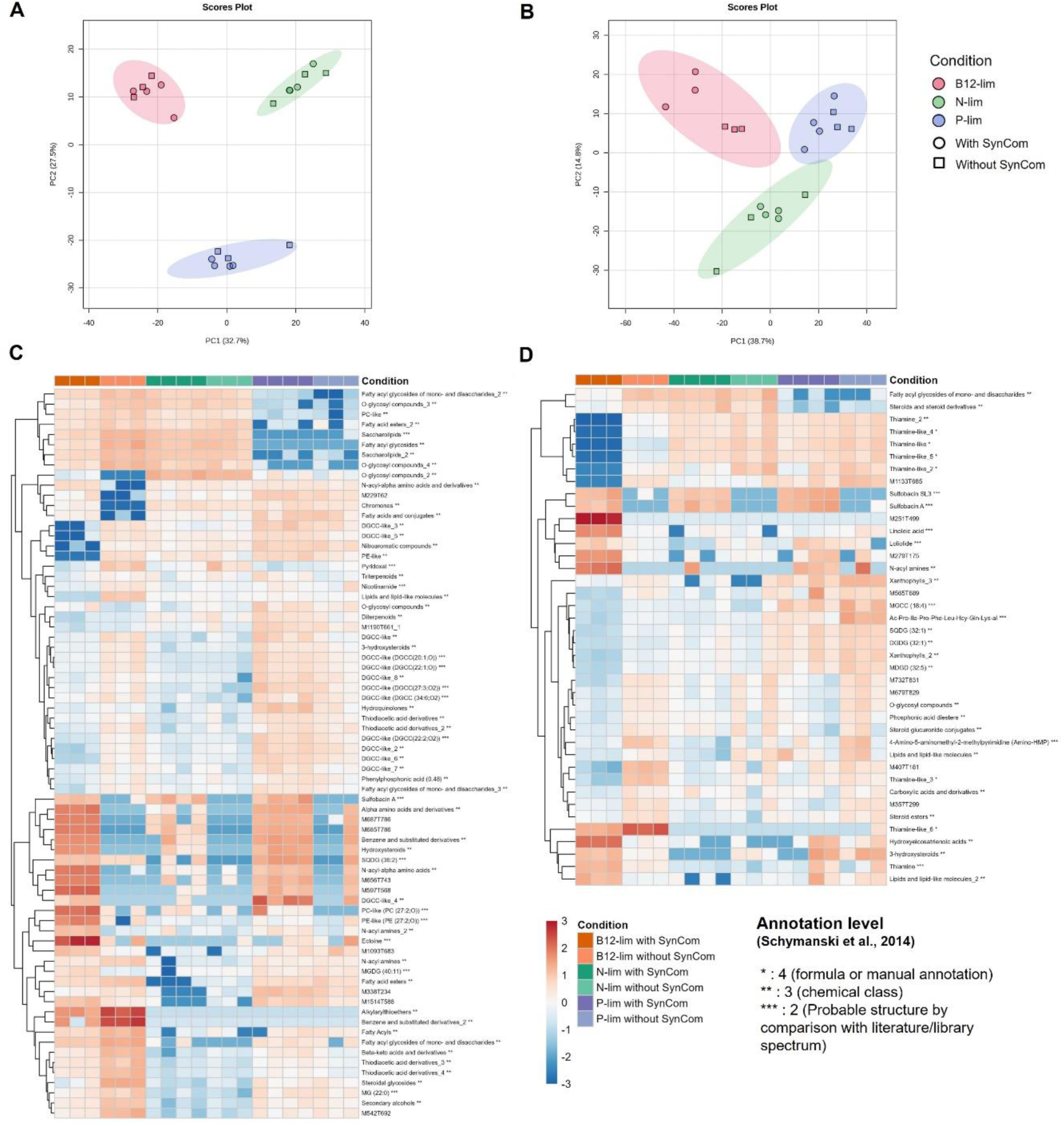
Multivariate and feature-level intra- and extracellular metabolomic responses to nutrient-limitation and SynCom presence. (A) Principal component analysis (PCA) of intracellular metabolites measured in *P. parvum* cultures under N-, P- and B12-lim, grown with or without the SynCom. Each point represents an individual sample, colored according to the nutrient-limitation condition and shaped according to SynCom presence or absence. (B) PCA of extracellular metabolites from the same cultures and conditions as in (A). (C) Heatmap of intracellular metabolic features significantly affected by the SynCom under B12-lim. Relative abundances of these features are shown across all nutrient-limited conditions (N-, P-, and B12-lim), with and without the SynCom, after normalization. Feature annotations indicate identification confidence levels following the classification proposed by Schymanski *et al.* (2014). (D) Heatmap of extracellular metabolic features significantly affected by the SynCom under B12-lim, displayed across all nutrient-limitation and SynCom treatment conditions as in (C).

### Metabarcoding analysis

The composition of the SynCom was followed over time and across nutrient limitations using 16S rRNA gene amplicon sequencing on both FL and PA fractions. A global NMDS analysis indicated that time (R² = 0.21, p < 0.001) was the strongest driver of variation, followed by fraction type (FL vs. PA; R² = 0.14, p < 0.001) and nutrient limitation (R² = 0.12, p < 0.001).

When considering each fraction separately, nutrient limitation primarily structured the FL community (R² = 0.20, p < 0.001), with time exerting a secondary influence (R² = 0.14, p < 0.001). In contrast, temporal variation dominated in the PA fraction (R² = 0.45, p < 0.001), while nutrient limitation contributed more modestly (R² = 0.11, p = 0.019; Figure 4A and 4B).

**Figure 4.**
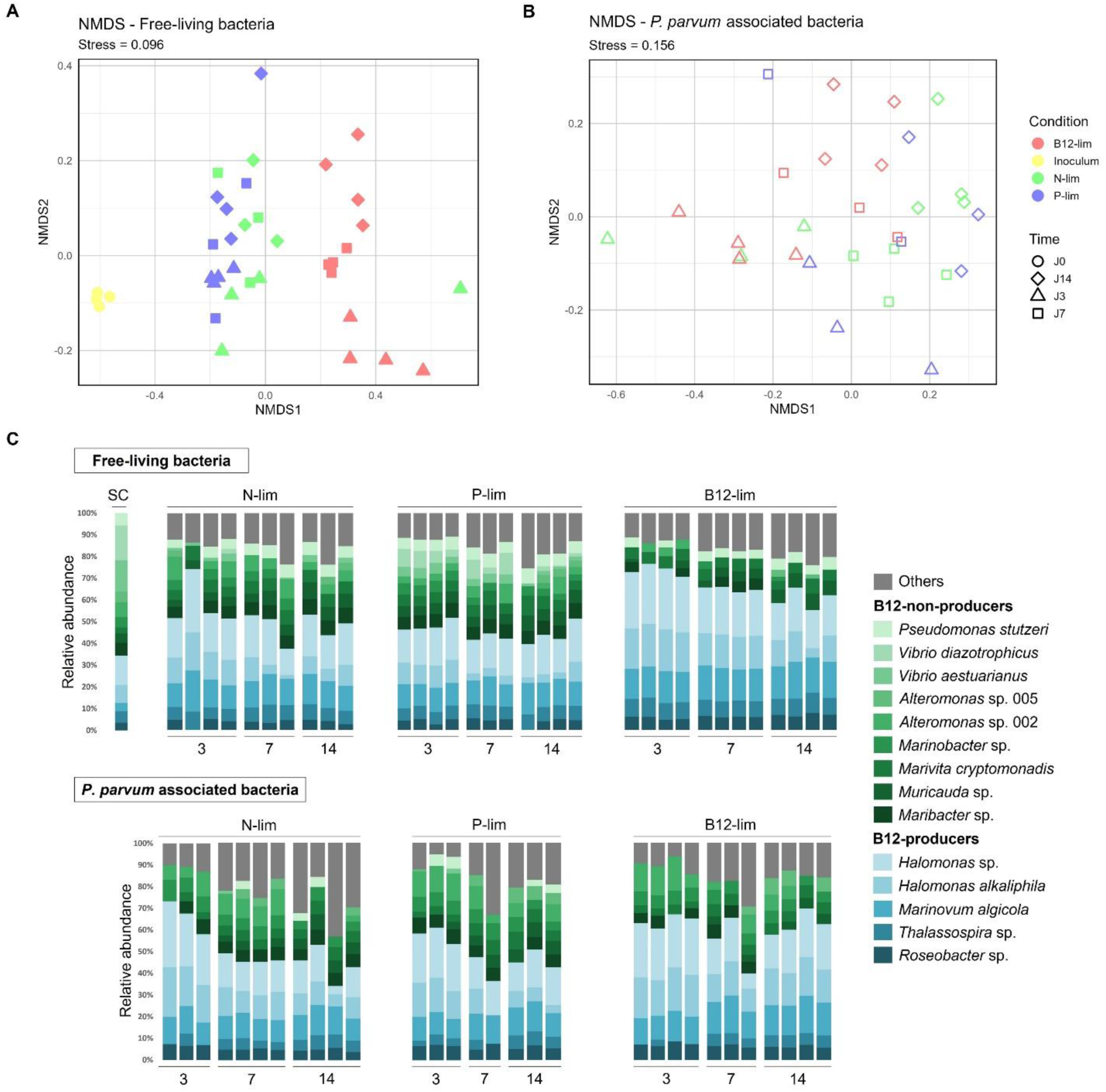
Dynamics of free-living and P. parvum-associated bacterial communities under nutrient limitation. (A) Non-metric multidimensional scaling (NMDS) ordination of Bray-Curtis dissimilarities based on 16S rRNA metabarcoding data from free-living (FL) bacterial communities, obtained by filtration through 3 µm and 0.2 µm filters. Points are colored according to nutrient-limitation condition (N-, P-, or B12-lim) and shaped according to sampling time (0, 3, 7, 14 days). (B) NMDS ordination of *P. parvum*-associated (PA) bacterial communities presented as in (A). (C) Relative abundance of SynCom bacterial taxa in FL (top) and PA (bottom) fractions across nutrient-limitation conditions and time points. Barplots show the 14 SynCom strains detected by 16S rRNA metabarcoding (with *Cyclobacterium qasimii* excluded as it was not detected), while all non-SynCom taxa are grouped as “Others” (grey). The “SC” bar represents the initial SynCom composition at T0 prior to inoculation. Taxa are colored by functial trait of interest, the production of vitamin B12: B12-non-producers in green and B12-producers in blue.

Across all conditions, fourteen of the fifteen SynCom strains were consistently detected. Due to its low abundance (below the rarity threshold), *Cyclobacterium qasimii* was removed during the bioinformatic decontamination pipeline and excluded from subsequent analyses. Only ASVs corresponding to SynCom members were retained, with all other sequences grouped as “Others” (Figure 4C).

Several SynCom taxa exhibited fraction- and condition-specific patterns. *Pseudomonas stutzeri* was enriched in the FL fraction across all conditions, whereas *Vibrio* strains occurred exclusively in the FL fraction under P-lim. Both *Alteromonas* strains, were specifically enriched in the PA fraction under B12- and P-lim. Finally, B12-producing bacteria (*Marinovum algicola*, *Thalassospira* sp., *Roseobacter* sp., *Halomonas alkaliphila*, and *Halomonas* sp.) were enriched in B12-lim cultures, which together reached 60% ± 4% relative abundance at day 14, compared to 46% ± 5% under N- and P-lim. These results show a globally stable SynCom despite nutrient variation with an enrichment in B12 producers under vitamin scarcity.

### Transcriptomic analysis

To investigate molecular mechanisms underlying differences in community structure and potential interactions between *P. parvum* and the SynCom under nutrient limitations, we analyzed day-14 transcriptomes for N- and P-lim with the SynCom, and for B12-lim with and without the SynCom. Although this represents a single time-point snapshot, it captures a stage where the systems have diverged substantially, providing valuable insights into the transcriptomic activities of both *P. parvum* and the bacterial consortium and into how nutrient availability shapes their interactions.

### Transcriptional responses of *P. parvum* to nutrient limitations in the presence of the SynCom

Analysis of *P. parvum* mRNA on day 14 revealed that 98.3% of genes were expressed at comparable levels across N-, P-, and B12-lim, while only 1.7% (443 genes) were differentially expressed (|log₂FC| > 3, padj < 0.05; Figure 5A, Table S3). DEGs were grouped into four clusters showing distinct functional enrichments (Figure 5B), although most (64-80%) lacked annotation.

**Figure 5.**
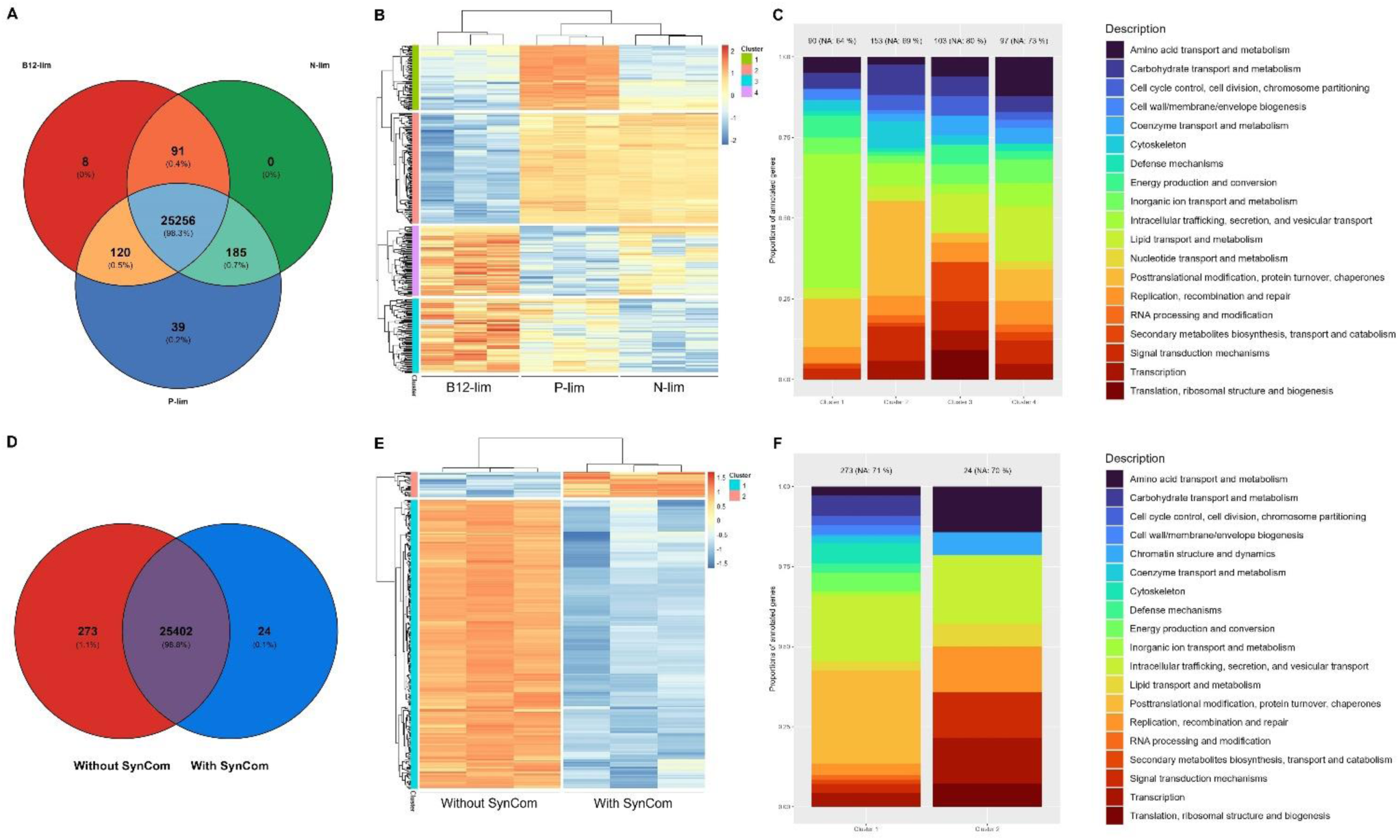
Transcriptional responses of P. parvum to nutrient limitation and SynCom presence. (A) Venn diagram showing the sets of *P. parvum* genes expressed under N-, P-, and B12-lim in the presence of the SynCom. (B) Heatmap of genes significantly differentially expressed across nutrient-limited conditions in the presence of the SynCom, clustered according to shared expression patterns. (C) Functional overview of transcriptional clusters identified in (B). Barplots summarize the main functional categories represented in each of the four clusters; bar colors correspond to functional groups. Numbers above bars indicate the total number of genes per cluster, with “NA: X” denoting the number of unannotated genes. (D) Venn diagram showing the sets of *P. parvum* genes expressed under B12-lim in the presence or absence of the SynCom. (E) Heatmap of genes significantly differentially expressed between B12-lim cultures with and without the SynCom, clustered based on shared expression profiles. (F) Functional overview of transcriptional clusters identified in (E). Barplots summarize the main functional categories represented in each of the two clusters; bar colors and annotations are as in (C).

Cluster 1 comprised 90 genes upregulated under P-lim but downregulated under N- and B12-lim, enriched in intracellular trafficking, secretion, and vesicular transport, with several photosynthesis-related genes and genes for phosphorus transport/remobilization, suggesting enhanced resource recycling to compensate for P limitation. Cluster 2 included 153 genes upregulated under P- and N-lim but repressed under B12-lim, enriched in posttranslational modification, protein turnover, and chaperone functions, including carbon metabolism and stress response genes which indicates activation of general stress programs under macronutrient limitation. Cluster 3 consisted of 103 genes upregulated under B12-lim and moderately under P-lim, but downregulated under N-lim, dominated by genes of unknown function plus transcription/translation-related genes, suggesting the activation of the growth machinery under B12-lim with SynCom. Cluster 4 comprised 97 genes upregulated under B12-lim and moderately under N-lim, enriched in amino acid and nucleotide transport/metabolism, including multiple genes for nitrogen metabolism and transport which suggest that nitrogen becomes the next liming factor once B12 limitation is alleviated (Figure 5C).

Of particular interest, the B12-dependent methionine synthase gene *metH* was expressed in all three nutrient limitation conditions without significant differences in expression suggesting an obligate dependency on B12 (Table S4).

### Transcriptional analysis of *P. parvum* under B12 limitation depending on the presence of the bacterial community

To explore the influence of the SynCom on *P. parvum* under B12 limitation, transcriptomes were compared on day 14. Most genes (98.8%) were expressed at comparable levels with or without the SynCom, while 1.2% were condition-specific, corresponding to 297 DEGs (|log₂FC| > 3, padj < 0.05; Figure 5D, Table S5) grouped into two clusters (Figure 5E). Cluster 1 (273 genes) was upregulated without the SynCom, enriched in posttranslational modification, protein turnover, and chaperones, suggesting a stress-related state. Cluster 2 (24 genes) was upregulated with the SynCom and repressed without, including genes involved in amino acid transport/metabolism, transcription, translation, replication, and unknown functions, signature of growth reactivation (Figure 5F).

The *metH* gene was expressed at similar levels in both conditions also confirming *P. parvum* B12 dependency (Table S4).

### Transcriptional analysis of SynCom in co-culture with *P. parvum* under different nutrient limiting conditions

A Venn diagram combining gene presence/absence and differential expression showed that 35.3% of bacterial genes were expressed across all nutrient conditions, 6.7% were shared between at least two conditions, and the remainder were condition-specific: 45.2% B12-lim, 11.2% N-lim, 1.5% P-lim (Figure 6A). Expression varied among SynCom members: *V. aestuarianus* showed no detectable expression, *V. diazotrophicus* a limited activity (280 genes), while other strains expressed 1,488 to 5,264 genes (Figure 6B).

**Figure 6.**
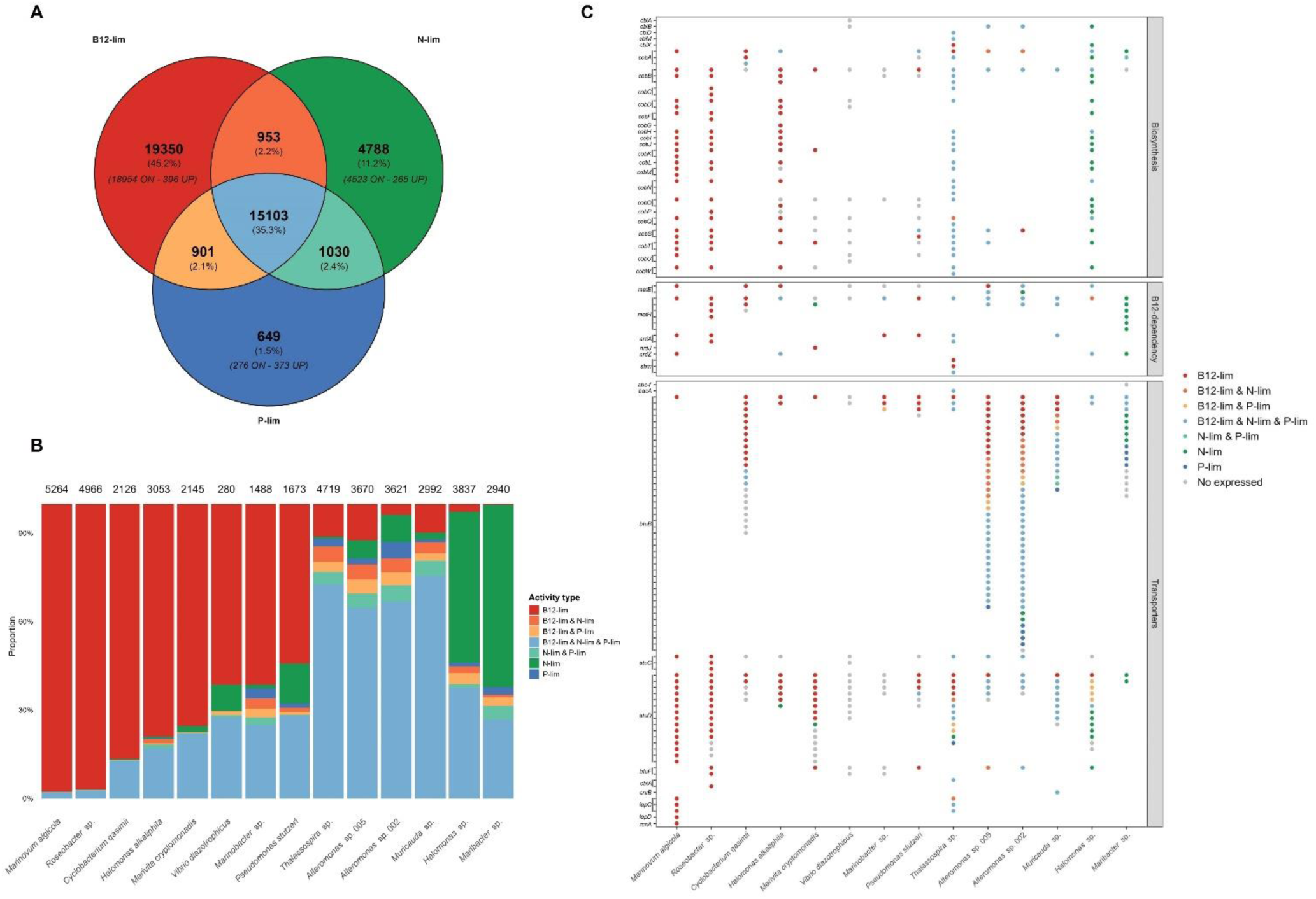
Transcriptional profiles of the SynCom under nutrient-limited conditions. (A) Venn diagram summarizing bacterial gene expression in *P. parvum* co-cultures under N-, P-, and B12-lim at day 14. Values in bold in each section indicate the total number of genes expressed and upregulated in the corresponding condition. Numbers below further distinguish genes that were upregulated (“UP”) from those expressed exclusively in a given condition (“ON”). (B) Distribution of expressed genes across SynCom bacterial strains. Barplots show the proportion of expressed genes for each of the 14 SynCom strains detected by metatranscriptomics (with *Vibrio aestuarianus* excluded as it was not detected). Numbers above bars indicate the total number of expressed genes per strain. Colors represent the nutrient-limitation conditions in which genes were expressed: single limitations (N-, P-, or B12-lim), pairwise combinations (N+P, N+B12, P+B12), or all three limitations (N+P+B12). (C) Expression patterns of targeted SynCom genes involved in vitamin B12 metabolism across nutrient-limiting conditions. The x-axis lists SynCom bacterial strains and the y-axis shows selected genes grouped into three functional categories: B12 biosynthesis genes, B12-dependent genes (including *metE* for comparison with *metH* expression), and known B12 transporter genes. Gene expression patterns are indicated by color.

Bacterial strains displayed four distinct patterns of expression. “Highly B12-lim-specific strains”, such as *Marinovum algicola* and *Roseobacter* sp., had over 90% of their genes uniquely expressed or upregulated under B12-limitation. Six “moderate B12-lim-specific strains”, including *Cyclobacterium qasimii* and *Halomonas alkaliphila*, had more than 50% of genes uniquely expressed or upregulated under B12-lim. “Generalist strains”, including *Thalassospira* sp., the two *Alteromonas* strains, and *Muricauda* sp., expressed over 60% of their genes across all three nutrient limitation conditions. Finally, “moderate N-lim-specific strains”, such as *Halomonas* sp. and *Maribacter* sp., had more than 50% of genes uniquely expressed or upregulated only under N-lim (Figure 6B). These strategies suggest complementary roles that maintain holobiont stability under fluctuating nutrient limitation.

COG enrichment analysis of the generalist bacteria revealed shared metabolic signatures across nutrient limitations (Figure S7). Under P-lim, *Alteromonas* sp. 002, *Muricauda* sp., and *Thalassospira* sp. increased functional potential related to inorganic ion transport and metabolism. Both *Alteromonas* strains upregulated genes associated to cell motility under N-and P-limitation, while transcription, replication, lipid metabolism, vesicular transport, and carbohydrate metabolism were predominantly upregulated under B12-lim by *Alteromonas* sp. 005, *Muricauda* sp., and *Thalassospira* sp. These patterns indicate that generalists provide flexible metabolic capacity that buffers the holobiont.

### B12 producers

Given that *P. parvum* resumed growth under B12-lim from day 9 only in the presence of the SynCom, we examined bacterial expression of vitamin B12-related genes across SynCom members to elucidate the contribution of each bacterial strain (Figure 6C). Five strains, *Marinovum algicola*, *Roseobacter* sp., *Halomonas alkaliphila*, *Halomonas* sp., and *Thalassospira* sp. possessed the complete B12 biosynthetic pathway. Among these, *Marinovum algicola*, *Roseobacte*r sp., and *Halomonas alkaliphila* exhibited a “B12-lim-specific” profile, with most genes, including those of the aerobic B12 biosynthetic pathway, expressed exclusively under B12-limited conditions, suggesting B12 production. *Thalassospira* sp. displayed a “generalist” pattern with broad expression of its B12 biosynthetic genes across all nutrient limitation conditions. Only three genes showed condition-specific regulation: *cbiM*, an essential component of the aerobic cobalamin biosynthetic pathway, and *cbiX*, whose annotation (the “X” denoting unknown function) prevents precise interpretation beyond its general implication in the pathway, were significantly upregulated under B12-lim. In addition, *cobQ*, which expression has been correlated to B12 production [53], was significantly upregulated under both B12- and N-lim. However, *cobQ* is not strictly essential, since its function can be complemented by *cbiP*, which catalyzes the same step in the pathway. *Halomonas* sp. showed a “N-lim-specific” profile, with most B12 biosynthetic genes upregulated under N-lim.

### B12 dependency

To assess B12 dependency within the SynCom, we analyzed the presence and expression of key B12-dependent genes. All 14 transcriptionally active strains possessed *metH*, encoding the B12-dependent methionine synthase, while nine strains also carried the B12-independent *metE*. Several strains contained additional B12-dependent genes (*nrdA, nrdJ, nrdZ, sbm*) (Figure 6C). Among non-B12 producers, *Cyclobacterium qasimii*, *Marivita cryptomonadis*, *Marinobacter* sp., and *Pseudomonas stutzeri* upregulated B12-dependent genes under B12-lim. Strains with both *metH* and *metE* showed differential regulation: *Marinovum algicola* and *Cyclobacterium qasimii* upregulated both, *Halomonas* sp., and *Pseudomonas stutzeri* upregulated *metH*, whereas *Halomonas alkaliphila* and *Alteromonas* sp. 005 upregulated *metE* under B12-lim.

### B12 transport

Finally, to investigate B12 potential transport, we identified known cobalamin transport genes across seven transporter families, including the canonical btu system (*btuB, btuC, btuD, btuF*; [54]; Figure 6C), implicated in B12 import. For the other transporter families, it remains unclear whether they mediate import or export of cobalamin. All bacterial members expect *Vibrio diazotrophicus* expressed B12 transporters under B12-lim, with the number of homologues varying among strains. Notably, the non-B12 producers *Cyclobacterium qasimii*, *Marivita cryptomonadis, Marinobacter* sp., and *Pseudomonas stutzeri*, which also upregulated B12-dependent genes, strongly induced most of their B12 transporters under B12-lim conditions. This widespread induction suggests active competition for extracellular B12 within the holobiont, coupled with efficient acquisition by dependent taxa.

## Discussion

By integrating phenotypic measurements with a multi-omics approach, this study provides new insights into *P. parvum*-bacteria interactions within the algal holobiont under contrasting nutrient limitations. We establish a link between algal physiology and bacterial strategies, revealing bidirectional interactions in which nutrient availability drives reciprocal modulation of algal and bacterial metabolism, highlighting the dynamic nature of the *P. parvum* holobiont under nutrient limitation.

### Mutualistic interactions within the holobiont are driven by nutrient limitations

Under N- and P-lim, co-culturing *P. parvum* with the SynCom did not significantly affect algal growth while under B12-lim, a strong mutualistic interaction emerged: algal growth increased more than three-fold in the presence of the SynCom, while bacterial abundances were also higher than under N- or P-lim. These reciprocal benefits are consistent with algal vitamin auxotrophy and bacterial provision of B12, supported by algal-derived resources.

Previous studies have shown that nutrient limitation can modulate algal-bacterial interactions by reshaping SynCom dynamics [55], and that associated bacteria can promote algal growth through vitamins and phytohormones [24]. Previous work further emphasized that metabolic context, particularly auxotrophy and nutrient stress, determines whether interactions result in mutualism, neutrality, or competition [56, 57]. Building on these findings, our study demonstrates that within our SynCom, vitamin B12 limitation constitutes the key context triggering strong reciprocal interactions with *P. parvum*, reinforcing the idea that metabolic dependencies such as vitamin auxotrophy are decisive drivers of holobiont mutualism [15, 58].

The absence of growth stimulation under N- or P-lim may reflect insufficient bacterial nutrient cycling or preferential retention of regenerated nutrients by bacteria. Although associated bacteria can recycle nutrients and supply vitamins such as B12 [59], such support may be insufficient or mismatched to algal demand. Preferential nutrient retention can further reduce availability to phytoplankton [60], and mutualistic interaction through nitrogen remineralization strongly depends on the physiological state of both partners [61]. In addition, for nitrogen fixation, the oxygen sensitivity of the nitrogenase complex may have constrained activity in our oxic batch cultures [62], suggesting that microscale organization [63] or protective respiration [64] may be required for non-cyanobacterial diazotrophs to sustain algal growth under aerobic conditions.

### Prymnesin production is induced by macronutrient stress

Beyond growth dynamics, nutrient limitation profoundly shapes specialized metabolism, notably toxin production. Phosphorus stress has previously been associated with enhanced toxicity in *P. parvum*, potentially reflecting a reallocation of cellular resources toward secondary metabolism when growth is constrained [65, 66] and with ecological consequences documented in natural blooms [67]. While earlier studies relied on indirect measurements, mass spectrometry analyses have confirmed that phosphorus limitation enhances prymnesin production [43].

Here, we provide the first direct comparison of prymnesin production under nitrogen, phosphorus, and vitamin B12 limitation at a similar growth phase. Our results show that prymnesin accumulation is specifically modulated by macronutrient stress rather than by micronutrient limitation, refining previous notions that nutrient stress broadly triggers toxin production [33, 68, 69]. One explanation is that N- and P-lim generate excess fixed carbon that can be redirected toward secondary metabolism, whereas B12-lim primarily impairs cofactor-dependent enzymatic reactions and slows overall metabolism [70], preventing carbon accumulation and toxin biosynthesis.

The presence of the SynCom did not significantly affect prymnesin levels under any nutrient condition, consistent with previous observations [36], but contrasting with other systems such as *Alexandrium pacificum* co-cultured with *Jannaschia cystaugens*, where bacterial presence significantly increases toxin production [71]. This suggests that in *P. parvum*, prymnesin biosynthesis seems tightly coupled to macronutrient stress, rather than being strongly modulated by bacterial cues, although other ecological factors may contribute [72].

### A potential holobiont metabolic signature of *P. parvum* resumed growth

Nutrient limitation reshaped both the endo- and exometabolomes of *P. parvum*, with a clear separation of N-, P-, and B12-lim conditions. Comparable stress-driven metabolic shifts have been reported in other algal groups [73]. Under B12-lim, metabolomic profiles further distinguished cultures with and without the SynCom, consistent with previous reports showing that microbial partners influence algal metabolic landscapes [57, 74, 75]. These findings highlight that metabolomic outputs integrate both endogenous adjustments to nutrient stress and exogenous modifications mediated by microbial associations.

Across all nutrient limitations, sulfobacin-like compounds were consistently enriched in co-cultures, suggesting a bacterial origin. Although sulfonolipids such as sulfobacins have not previously been described in algal-bacterial systems, their structural similarity to bioactive sulfonamides which exhibit antibacterial activity [76, 77], raises the possibility that they could contribute to microbial interactions or pathogen defense.

Under B12-lim, co-cultures showed a distinct metabolic profile characterized by ectoine accumulation, consistent with enhanced osmoprotection [78]. Ectoine, produced primarily by halophilic bacteria such as *Halomonas* spp., and also reported in *P. parvum* [78, 79], protects cells against multiple stressors [80] and may reflect a joint survival strategy enhancing holobiont resilience. In parallel, increases in membrane lipids suggest membrane-remodeling [81] preceding growth recovery. The function of the strongly accumulated unknown metabolite M251T499 remains unclear. Some metabolites enriched in co-culture, including ectoine and PC/PE lipids, may also provide carbon and nitrogen substrates for bacteria [82, 83]. Conversely, thiamine-like compounds were strongly reduced, consistent with cofactor reallocation under B12-lim [84]. Decreases in glycerolipids, glycosylated metabolites, and DGCC-like lipids further suggest attenuation of stress-related lipid pools [85]. Although cultures remained in plateau phase at day 8, these metabolic shifts anticipated the growth recovery observed at day 10, indicating that the SynCom facilitated a transition toward renewed algal proliferation.

### Transcriptional reprogramming of *P. parvum* under nutrient limitation

Transcriptomic profiling revealed nutrient-specific transcriptional responses in *P. parvum*. Under P-lim, responses appear to extend beyond phosphate acquisition to include vesicular trafficking and photosynthesis-related functions, suggesting broad cellular reorganization to optimize resource use. N-lim was characterized by induction of stress-related genes alongside nitrogen transporters and metabolic enzymes, indicating concurrent activation of coping mechanisms and assimilation pathways. In both conditions, stress-responsive transcripts predominated, consistent with the possibility of a generalized stress program involving protein turnover, chaperone activity, and metabolic downscaling as previously described [68, 86]. By comparing cultures at the same physiological state under well-defined N, P, and B12 limitation, our study isolates nutrient-specific responses while minimizing growth-stage effects.

In contrast, under B12-lim, the presence of the SynCom induced a distinct trajectory, with a shift toward growth-associated gene expression, including transcription, translation, amino acid metabolism, and nitrogen transport. This pattern indicates that bacterial B12 provisioning enabled *P. parvum* to resume metabolic activity, with nitrogen emerging as the next limiting resource. Similar vitamin-mediated recoveries have been documented in other algal-bacterial systems [15, 19, 58], underscoring the general importance of vitamin exchange in shaping algal physiology.

### B12 fluxes within *P. parvum* holobiont

Metatranscriptomic profiling of the SynCom revealed functional specialization underlying holobiont-level responses. Although community composition remained broadly stable, with only a slight enrichment of B12 producers under B12-lim, metatranscriptomics revealed contrasting bacterial strategies, with condition-specific specialists (e.g., B12 producers under B12 lim) and broadly responsive generalists contributing to community function, consistent with recent ecological frameworks [87, 88]. Enrichment of B12-producing taxa (*i.e*., *Marinovum algicola*, *Roseobacter sp.*, *Halomonas sp.*) under B12-lim coincided with the upregulation of cobalamin biosynthesis pathways, enabling algal growth recovery and induction of growth-associated algal genes. Similar bacterial B12 provisioning has been shown to restore algal metabolism in other systems [58, 89, 90], although such exchanges are not universal [19]. Together, metabarcoding and metatranscriptomics show that SynCom resilience stems not only from membership but also from functional reprogramming under nutrient stress.

SynCom bacteria displayed complementary B12-related strategies along three axes: production, dependence, and import (Figure 7). B12-prototrophs expressed the biosynthetic pathway, whereas auxotrophs relied on external sources. B12-dependent taxa expressed only *metH*, while independent taxa expressed *metE* and *metH*, allowing methionine synthesis regardless of vitamin availability. Notably, B12 independence does not preclude auxotrophy, as several non-producing taxa expressed *metE* while lacking the biosynthetic pathway. All members expressed B12 transporters, which were strongly upregulated under B12 limitation alongside B12-dependent enzymes, reflecting active acquisition and reinforcing functional complementarity.

**Figure 7.**
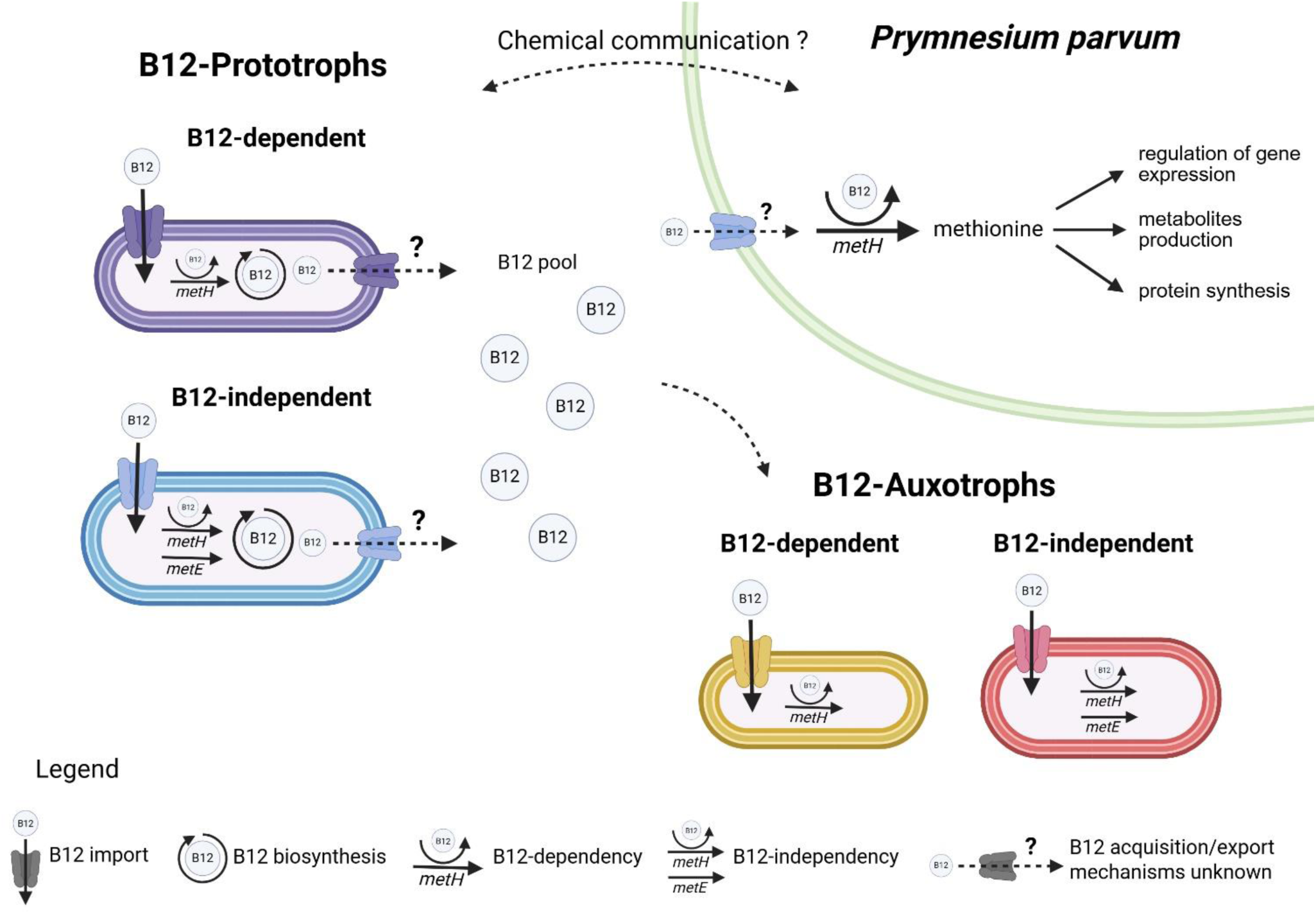
Conceptual model of vitamin B12-mediated interactions within the algal-bacterial holobiont. Bacterial strains are classified according to the expression of genes involved in vitamin B12 biosynthesis and B12 dependence based on methionine synthesis pathways. B12-dependent bacteria rely exclusively on the B12-dependent methionine synthase (*metH*), whereas B12-independent bacteria express both *metE* and *metH*. B12-prototrophs express the cobalamin biosynthetic pathway, while B12-auxotrophs lack its expression and rely on external sources of vitamin B12. All bacterial groups are able to import vitamin B12, while B12 release by producers is hypothesized and remains unresolved. Vitamin B12 exchanged within the holobiont supports methionine synthesis in the B12-dependent microalga *Prymnesium parvum*, thereby sustaining cellular growth and metabolism. Dashed arrows indicate hypothesized interactions. Created in BioRender. Réveillon, D. (2026) https://BioRender.com/om58389

The genomic features of *P. parvum* further shape this exchange. Like other algae that have evolved in close association with B12-producing bacteria [91], *P. parvum* lacks the B12-independent methionine synthase *metE* and relies exclusively on the B12-dependent isoform *metH*. Consistent with this obligate auxotrophy, *metH* was robustly expressed across all conditions and did not differ between cultures with or without the SynCom. This constitutive expression aligns with observations that microalgae do not modulate *metH* expression in response to intracellular B12 availability [92], maintaining the methionine cycle poised to exploit any available vitamin.

Together, these results suggest a holobiont-level mutualism. Heterotrophic bacteria dependent on algal organic matter synthesize and mobilize B12 under nutrient stress, and both partners utilize it. Such reciprocal adjustments exemplify mutualistic interactions consistent with models of symbiotic vitamin exchange in natural phytoplankton-associated communities [93].

The precise mechanism of B12 transfer remains unresolved. Not only are the potential acquisition routes unclear, but the identity of bacterial exporters mediating vitamin release is also unknown. Several possibilities are plausible, including uptake of extracellular vitamin passively released by bacteria [94], bacterivory [95], access to intracellular pools following bacterial lysis [96, 97], or even phage-mediated release, as recently shown in *Sulfitobacter* sp. M39 [98]. Although extracellular B12 was not detected, likely due to methodological constraints and low concentrations, the strong upregulation of bacterial importers and B12-dependent enzymes indicates that vitamin was available and actively utilized. These observations strongly suggest that passive release by producers and active uptake by dependent partners are central components of B12 exchange within the holobiont.

## Acknowledgements

We thank Prof. Alison Smith and Sue Aspinall (Department of Plant Sciences, University of Cambridge, UK) for providing the *Halomonas* sp. strain 19-001. We are grateful to François Delavat (US2B, Nantes Université, France) for providing the *Vibrio diazotrophicus* strain 23-002. We also thank Cyril Noël and the Sebimer platform (Ifremer, France) for their assistance with bioinformatics analyses, and Gregory Carrier (Phytox, Ifremer, France) for his support during genome sequencing. Finally, we acknowledge Jean-Baptiste Bérard (Phytox, Ifremer, France) for his help with the phenotyping bench during the initiation of the experiment.

