## Supplementary data for "Multi-omics uncovers nutrient stress-driven interactions in the *Prymnesium parvum* holobiont, with vitamin B12 limitation highlighting mutualism"

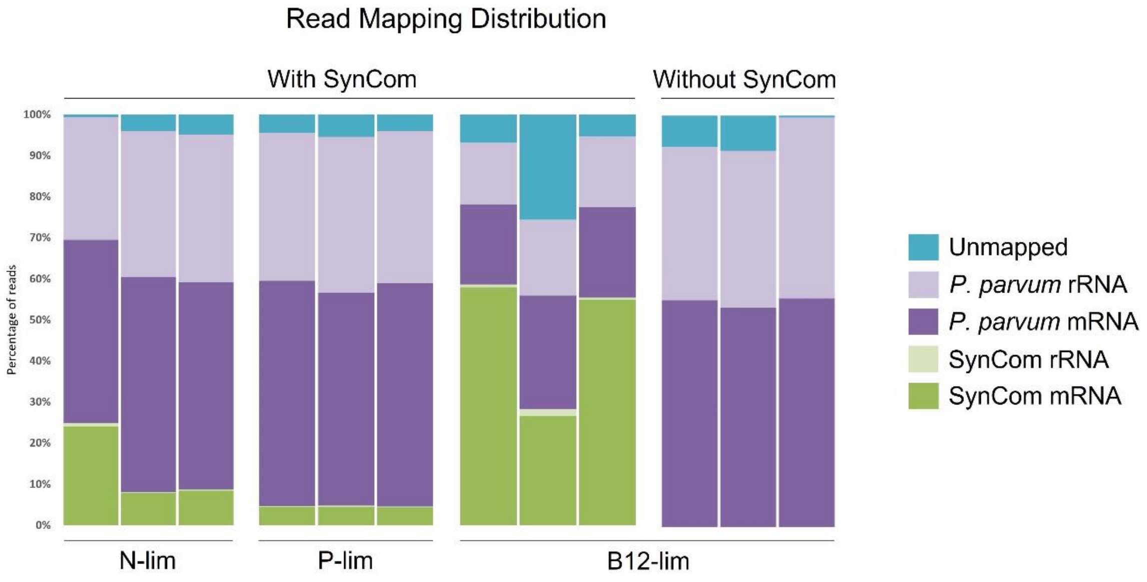


*Supplementary Figure S1. Distribution of transcriptomic reads across categories under N-, P-, and B12-lim on day 14.*

Each sample generated 30-40 million reads. In SynCom-containing cultures, read distribution varied according to nutrient condition: *P. parvum* mRNA accounted for 42 ± 15% of reads, *P. parvum* rRNA for 29 ± 10%, SynCom rRNA for 1 ± 1%, and unmapped reads for 6 ± 8%. SynCom mRNA abundance differed markedly across conditions, representing 47 ± 17% of total reads under B12-lim but only 9 ± 8% under N- and P-lim. In *P. parvum* monocultures, reads mapped predominantly to *P. parvum*: 55 ± 1% to mRNA and 40 ± 4% to rRNA, with unmapped reads contributing 6 ± 4% of the total.

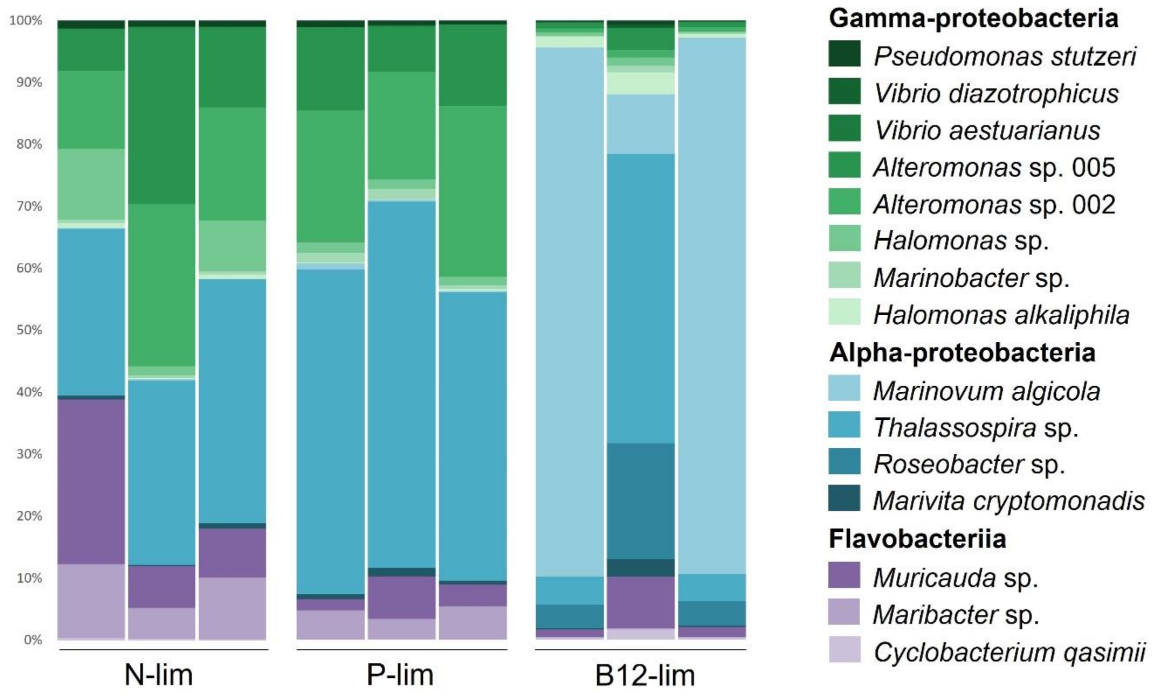


*Supplementary Figure S2. Taxonomic composition of SynCom mRNA reads under N-, P-, and B12-limitation on day 14.*

Under N- and P-lim, *Thalassospira* sp. dominated the SynCom transcriptome (42% ± 13% of reads), with the two *Alteromonas* strains contributing 34% ± 13%. In B12-lim, *Marinovum algicola* and *Roseobacter* sp. became dominant (61% ± 44% and 9% ± 9%), while other strains, including the two *Vibrio* spp., *Cyclobacterium qasimii*, and *Marinobacter* sp., each contributed less than 2%.

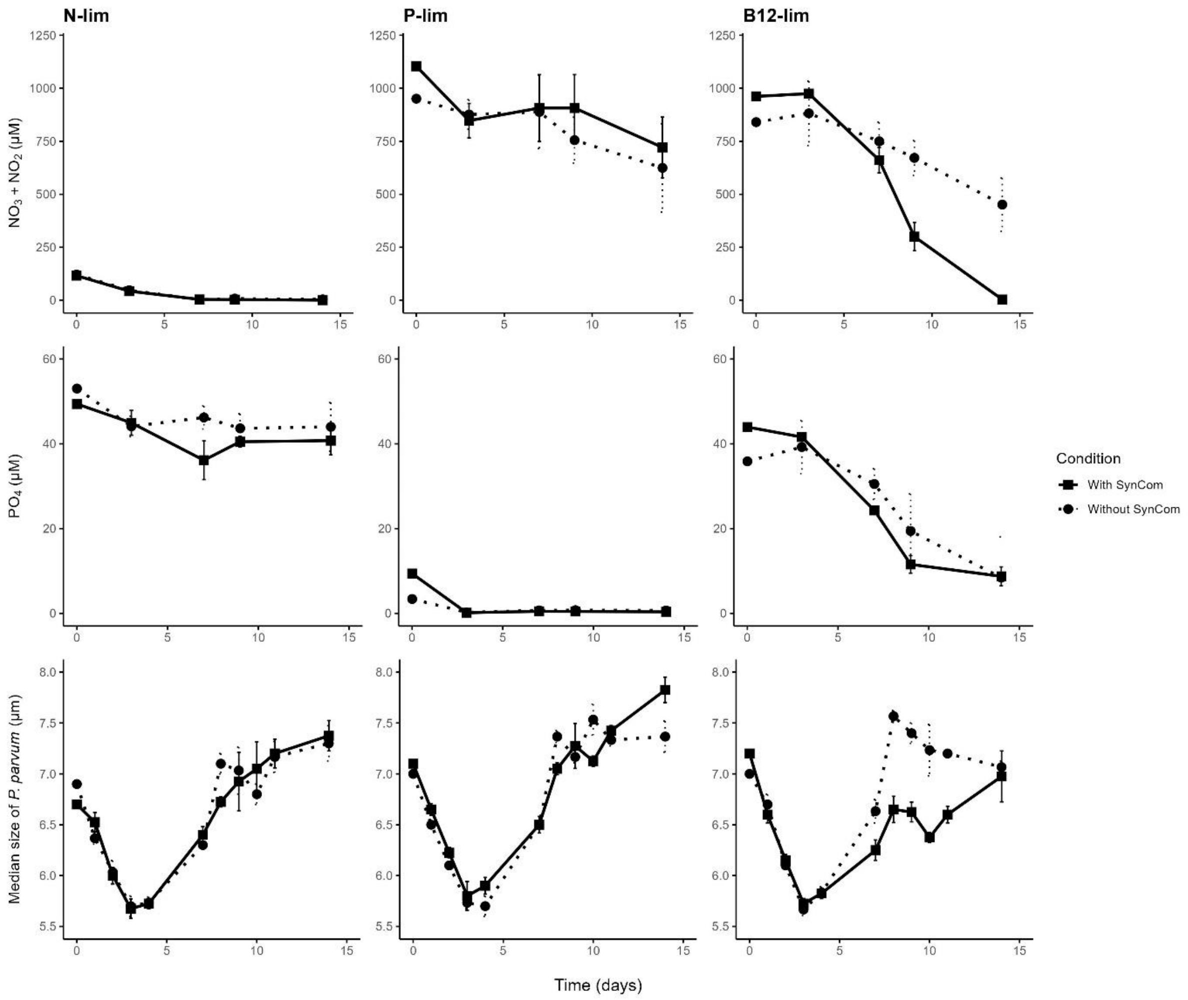


*Supplementary Figure S3. Individual growth curves of* P. parvum *with and without SynCom.*

(A) Growth curves of *P. parvum* in the presence of the SynCom. Each plot shows cell concentration (cells/mL) over time for one biological replicate. Plots are arranged in a 3 × 4 grid: rows indicate nutrient limitation (Row 1: N-lim, Row 2: P-lim, Row 3: B12-lim), and columns indicate spike type applied at day 14 (Column 1: N spike, Column 2: P spike, Column 3: B12 spike, Column 4: Seawater control). Each subplot corresponds to one replicate (n = 4 per condition), labeled at the top ("Replicate 1", etc.). The red dashed vertical line indicates the spike day (day 14). (B) Growth curves of *P. parvum* in the absence of the SynCom. Plots are arranged in a 3 × 3 grid (one replicate missing), with rows for nutrient limitation (N-lim, P-lim, B12-lim) and columns for spike type (N, P, B12).

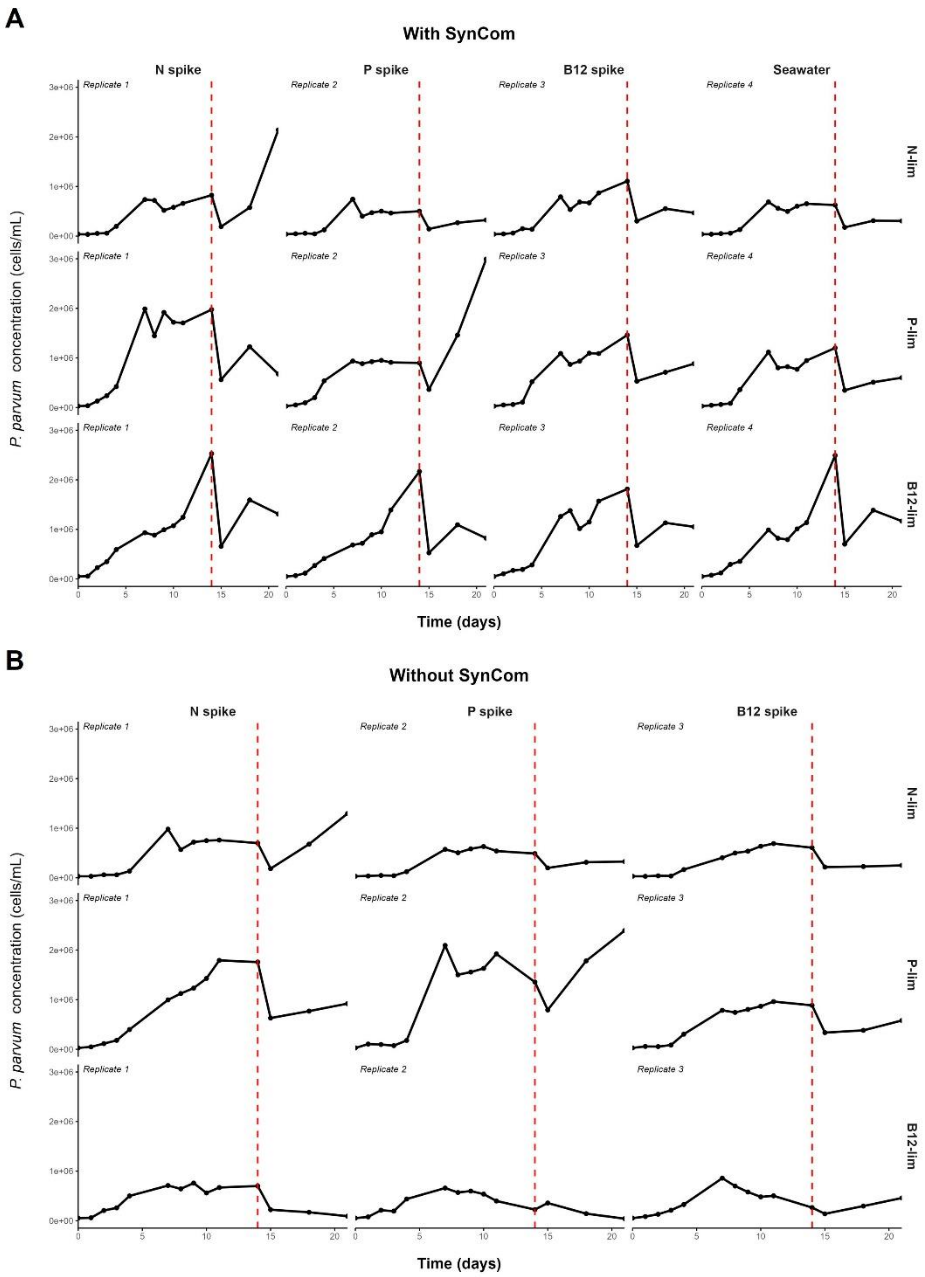


*Supplementary Figure S4. Temporal dynamics of physiological and environmental parameters under nutrient-limited conditions.*

Each column represents a nutrient limitation regime: N-lim; P-lim and B12-lim cultures. From top to bottom, the panels show: Dissolved inorganic nitrogen ($NO_3^-$ + $NO_2^-$, in µM), Dissolved phosphate ($PO_4^{3-}$, in µM), and median cell size of *P. parvum* (in µm). Data are plotted over time. Solid lines correspond to the With SynCom condition, and dashed lines represent the Without SynCom condition. The median cell size was acquired using a Multisizer 4 Coulter Counter (Beckman Coulter, Indiana, USA).

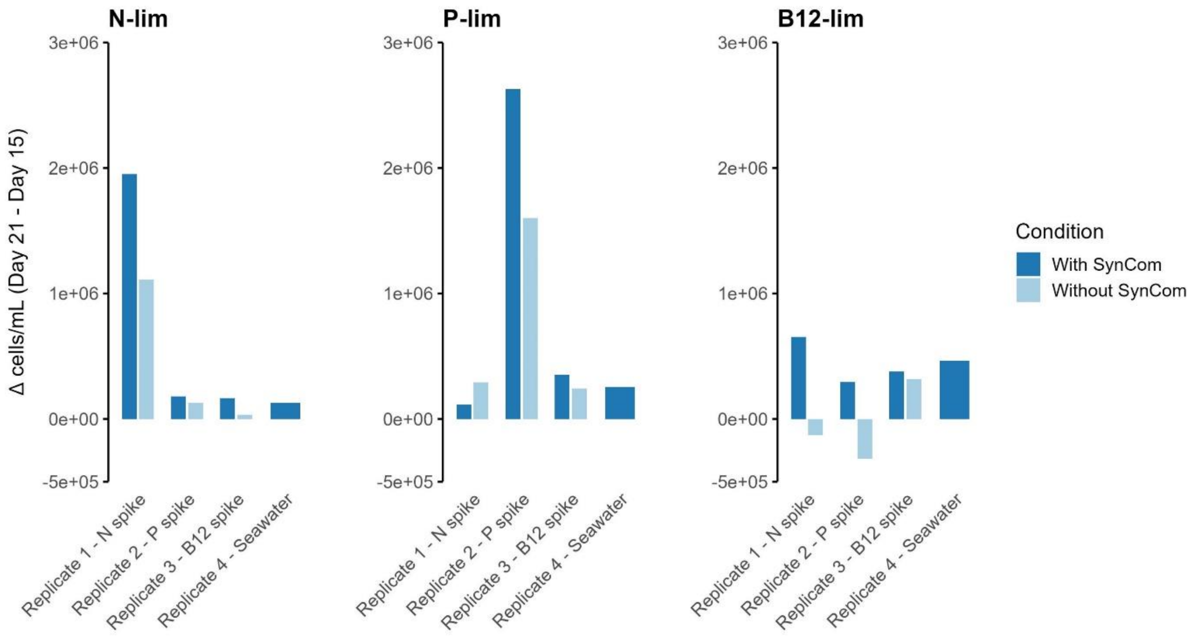


*Supplementary Figure S5. Effect of nutrient spikes on P. parvum growth recovery.*

Barplots show the change in algal cell concentration (Δ cells/mL between day 21 and day 15) under three limitation regimes: (A) N-lim, (B) P-lim, and (C) B12-lim cultures. Each plot displays four replicates corresponding to different nutrient spikes (N, P, B12, or seawater control) as well as the presence or not of the SynCom.

In N- and P-lim conditions, only the addition of the corresponding nutrient restored growth. Under B12-lim, all spikes induced moderate growth increases in the presence of SynCom, whereas without SynCom only B12 addition produced a response. Depletion of dissolved N and P (Figure S2), as well as elevated C/N ratios in N-lim cultures (Figure S4), further validated the limitation regimes.

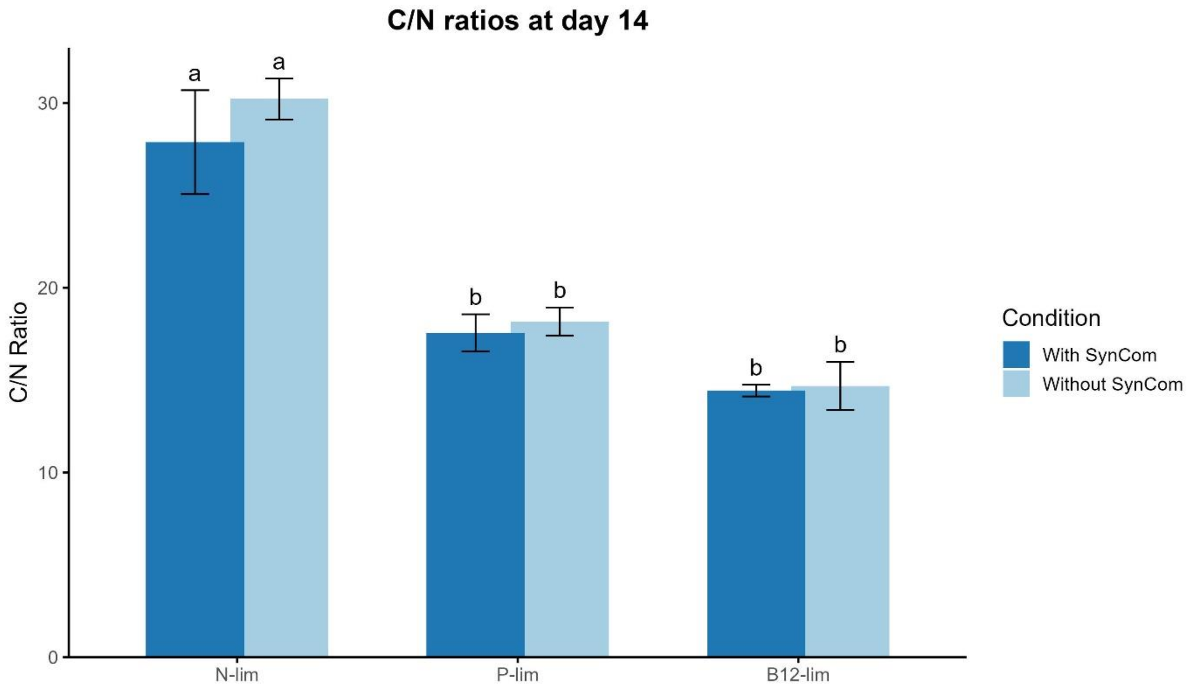


*Supplementary Figure S6. C/N ratios of P. parvum at day 14 under nutrient limitation.*

The barplot shows C/N ratios measured in cultures limited in N-, P- and B12-lim conditions at day 14 with or without the SynCom.

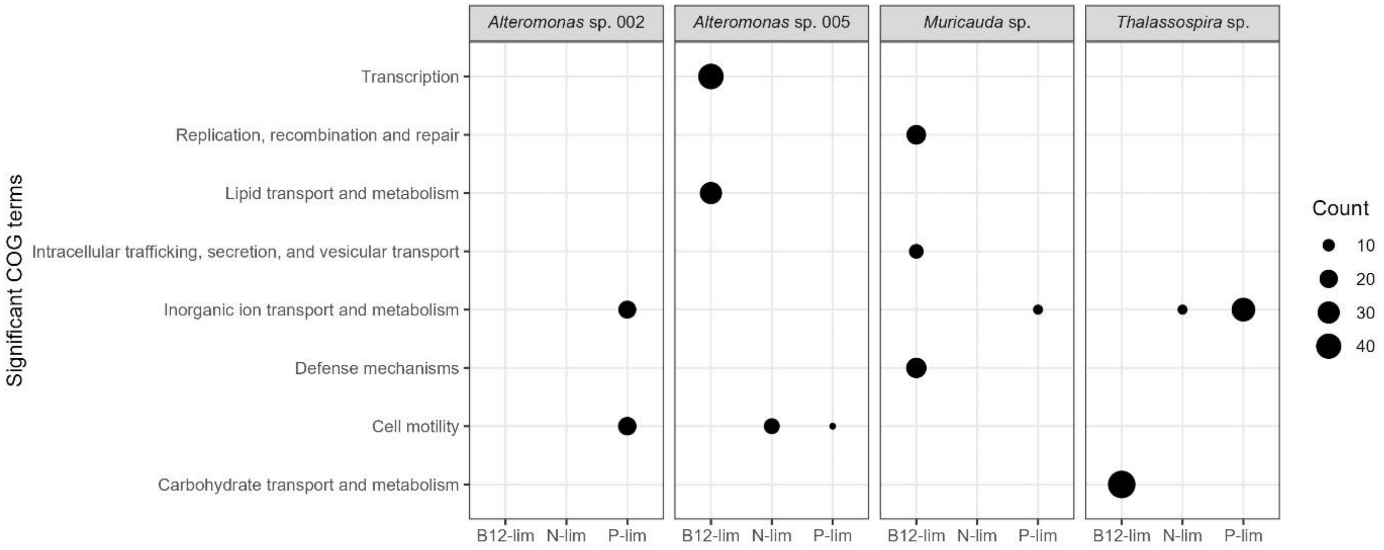


*Supplementary Figure S7. Functional enrichment analysis of the four plastic-responsive bacterial strains.*

Each panel corresponds to one of the four plastic-responsive bacterial strains. Within each panel, the X-axis shows the nutrient-limitation conditions (B12-, N- and P-lim), and the Y-axis lists the significantly enriched COG functional categories. Circles represent COG categories in which genes were significantly overexpressed under the corresponding nutrient-limitation condition; circle size is proportional to the number of overexpressed genes in that category.

*Supplementary Table S1. Composition of the SynCom and access to the genomes.*

*Supplementary Table S2. Metabolomics analysis of intracellular and extracellular metabolites significantly affected by the presence of the SynCom.*

This table lists metabolites that were significantly differentially produced between conditions with and without the SynCom, for both intracellular (Table S2A) and extracellular (Table S2B) fractions. Each row corresponds to a metabolomic feature identified by its unique MXXTXX code. Reported parameters include fold change (FC), log2(FC), adjusted p-value (p.adjusted), -log10(p), and significance status, as well as mass-to-charge ratio (m/z), retention time (RT), adduct and isotope information, and MS/MS availability. Putative metabolite annotations were obtained using Flash Entropy, GNPS spectral library matching, and SIRIUS, including identity or cosine scores, shared peaks, molecular formulae, compound class predictions (CANOPUS), and annotation confidence levels according to the Schymanski et al. (2014) classification. Additional comments and detailed annotation information are provided where applicable.

*Supplementary Table S3. Differential gene expression analysis of Phaeodactylum parvum in response to the syncom under nutrient limitation.*

This table reports the 443 DEGs identified in *P. parvum* grown with or without the syncom under N-, P-, and B12-lim on day 14. Each row corresponds to one DEG, and columns include the comparison performed, gene identifier (GeneID), base mean expression (baseMean), log2 fold change (log2FoldChange), adjusted p-value (padj), and regulation status. Functional annotation fields comprise orthology assignments (seed_ortholog, eggNOG_OGs), annotation confidence levels, COG functional categories, gene descriptions and preferred names, as well as Gene Ontology (GO), enzyme commission (EC), and KEGG annotations (KO, pathways, modules, reactions, and related classifications). Additional annotations include BRITE hierarchies, transporters (KEGG_TC), carbohydrate-active enzymes (CAZy), metabolic reactions (BiGG), protein domains (PFAMs), clustering information, and sequence data. Remarks are provided where relevant.

*Supplementary Table S4. Differential expression analysis of the metH gene in Prymnesium parvum.* This table focuses on the expression patterns of the B12-dependant methionine synthase gene (*metH*) in *P. parvum*. Differential expression analyses include comparisons between nutrient-limitation conditions (N-, P-, and B12-lim) as well as comparisons between cultures grown with and without the syncom, under B12-lim. Reported values include gene identifier (GeneID), comparison performed, mean normalized expression (baseMean), log2 fold change (log2FoldChange), standard error of the log2 fold change (lfcSE), test statistic (stat), raw p-value (pvalue), and adjusted p-value (padj).

*Supplementary Table S5. Differential gene expression analysis of Prymnesium parvum under vitamin B12 limitation in response to the syncom at day 14.*

This table reports the 297 differentially expressed genes (DEGs) identified in *P. parvum* grown under B12-lim with or without the SynCom at day 14. Each row corresponds to one DEG, with columns indicating the comparison performed, mean normalized expression (baseMean), log2 fold change (log2FoldChange), adjusted p-value (padj), regulation status, and clustering assignment. Functional annotations include orthology information (seed_ortholog, eggNOG_OGs), annotation confidence level, COG functional categories, gene descriptions and preferred names, as well as Gene Ontology (GO), enzyme commission (EC), and KEGG annotations (KO, pathways, modules, reactions, rclass, BRITE hierarchies, transporters, CAZy families, and BiGG reactions). Protein domain information (PFAMs) and sequence data are also provided.
